# Trichomes on female reproductive tract: rapid diversification and underlying gene regulatory network in *Drosophila suzukii* and its related species

**DOI:** 10.1101/2022.01.21.477200

**Authors:** Kentaro M. Tanaka, Kanoko Takahashi, Gavin Rice, Mark Rebeiz, Yoshitaka Kamimura, Aya Takahashi

## Abstract

**Background:** The ovipositors of some insects are external female genitalia, which have their primary function to deliver eggs. *Drosophila suzukii* and its sibling species *D. subpulchrella* are known to have acquired highly sclerotized and enlarged ovipositors upon their shifts in oviposition sites from rotting to ripening fruits. Inside the ovipositor plates, there are scale-like polarized protrusions termed “oviprovector scales” that are likely to aid the mechanical movement of the eggs. The size and spatial distribution of the scales need to be rearranged following the divergence of the ovipositors. In this study, we examined the features of the oviprovector scales in *D. suzukii* and its closely related species. We also investigated whether the scales are single-cell protrusions comprised of F-actin under the same conserved gene regulatory network as the well-characterized trichomes on the larval cuticular surface.

**Results:** The oviprovector scales of *D. suzukii* and *D. subpulchrella* were distinct in size and spatial arrangement compared to those of a closely related species *D. biarmipes* and *D. melanogaster*. The comparisons of the size of the scales suggested that the apical cell area of the oviprovector has expanded upon the elongation of the ovipositor plates in these species. Our transcriptome analysis revealed that 43 out of the 46 genes known to be involved in the trichome gene regulatory network are expressed in the developing female genitalia of *D. suzukii* and *D. subpulchrella*. An antibody staining depicted the presence of Shavenbaby (Svb) in the inner cavity of the developing ovipositors of *D. melanogaster* at 44–48 h after puparium formation (APF). Also, *shavenoid* (*sha*) was expressed in the corresponding patterns in the developing ovipositors and showed differential expression levels between *D. suzukii* and *D. subpulchrella* at 48 h APF.

**Conclusions:** The oviprovector scales have divergent size and spatial arrangements among species. Therefore, these scales may represent a rapidly diversifying morphological trait of the female reproductive tract reflecting ecological contexts. Furthermore, our results showed that the gene regulatory network underlying trichome formation is adopted to develop the rapidly evolving trichomes on the oviprovectors of these flies.

## Background

The morphology of male external genitalia has been known to diverse rapidly among species and for this reason, it has been used for practical purposes as one of the taxonomic characters for species identification [1–3]. However, recent studies have revealed some cases of rapidly diversifying female genital traits [4–6]. In particular, the ovipositors of some insects, including Drosophilidae, are involved in genital coupling during copulation as external genitalia with the primary function to deliver eggs onto the food source for the emerging larvae [7, 8]. Thus, the genital structures involved in egg deposition are constantly under selective pressure to fulfill this function and can experience rapid morphological evolution upon occasional shifts in oviposition sites.

Such cases are well-documented in the species of Hawaiian *Drosophila* and *Scaptomyza*, which utilize a wide range of host plants in which their larvae grow [9, 10]. As predicted, a large diversity of the size and shape of the ovipositors have evolved in these species [11, 12]. In addition, another type of diverse trait in the internal structures of these ovipositors is noticeable. On the surface of the oviprovector (the membrane surrounding the vulva), there are scale-like polarized protrusions [12], termed as the “oviprovector scale” [13]. Although the functional role of these scales is not clear, it is likely to aid the transport and delivery of eggs onto the oviposition substrate [14]. Thus, the spatial arrangement of the scales should change following the morphological divergence of the ovipositors to support the mechanical movement of the eggs effectively. Therefore, these scales are likely to have gone through frequent rearrangement and may represent a rapidly diversifying morphological trait of the female genital organ.

The shape of the observed oviprovector scales resembles the well-studied trichomes on the larval cuticular surface [15–17]. Similar trichomes are found on various parts of the body cuticle and wings of Drosophilid flies; however, those on the oviprovector have been overlooked. The gene regulatory network underlying trichome formation is well-characterized by intensive studies on the larval denticles (reviewed in [18]). If the oviprovector scales are single-cell protrusions comprised of F-actin like other epidermal trichomes, it is intriguing to examine whether the same conserved gene regulatory network has been adopted.

Among species of non-Hawaiian *Drosophila*, *D. suzukii* and its sibling species *D. subpulchrella* are known to have acquired highly sclerotized and enlarged ovipositors with stiff bristles lined up on the edge upon their shifts in oviposition sites from rotting to ripening fruits [8, 19, 20]. Therefore, it would be insightful to depict how the characteristics of the oviprovector scales have changed in these drastically elongated ovipositors. Moreover, if the trichome-producing gene regulatory network is involved, there should be changes in the expression patterns of some genes in the network associated with the differential scale arrangements.

A recent study on the developing pupal ovipositors of *D. suzukii* and *D. melanogaster* showed that the timing of cell proliferation and the final cell number are similar between the two species [21]. By tracking the cell shape, the authors conclude that the expansion of the apical area of the cells is responsible for the development of the elongated ovipositor plates (hypogynium valves) in *D. suzukii*. Given these observations on the surface cell layer of the ovipositor plates, we can ask whether the cell size expansion has also taken place in the simultaneously elongated oviprovector in this species.

In this study, we first show the divergence of the size and distribution of the oviprovector scales in *D. suzukii* and *D. subpulchrella* and their closely related species, *D. biarmipes* and *D. melanogaster*. Next, we indicate that the oviprovector scales are single-cell protrusions like the trichomes developed on the larval cuticle and provide inferences about the cell size changes of the egg duct membrane. Finally, we report that an effector gene, *shavenoid* (*sha*), involved in the actin-bundling during the trichome development, is differentially expressed between the developing ovipositors of the two species, resembling the locations where the trichomes align in the later stages. From these results, we aim to emphasize the importance of the diversifying inner ovipositor structures in response to the rapid ovipositor shape adaptation and report for the first time that a trichome network gene is involved in the development of the divergent oviprovector scales.

## Results

### Divergent oviprovector scales

The oviprovector scales reside on the inner surface of the oviprovector (egg duct) in *Drosophila* species [13] and presumably aid egg deposition (Fig. 1A, B). The SEM image of the everted oviprovector of *D. suzukii* depicted the scales covering the inner surface of the duct (Fig. 1C, 1C’) as in similar SEM images shown previously by Hamby et al. [22]. Since the scales are aligned parallel to the movement of the eggs through the duct and are likely to guide the eggs upon oviposition, they may have been rearranged during the rapid elongation of the ovipositors in *D. suzukii* and *D. subpulchrella*.

**Fig. 1.**
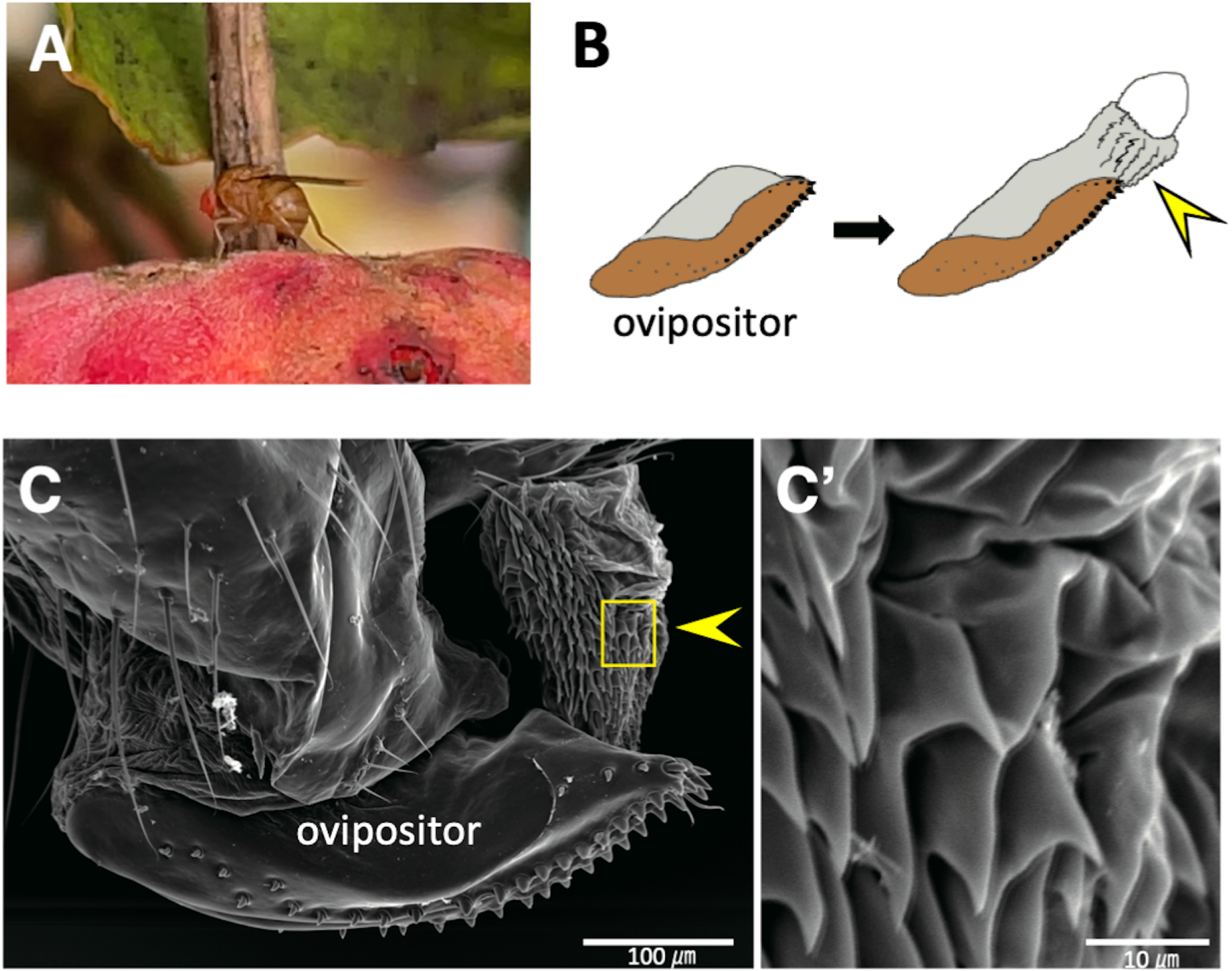
Oviprovector scales of *Drosophila suzukii*. (A) Adult female laying an egg into a fruit of *Cornus kousa*. (B) Oviprovector scales on the reverted membrane (indicated by an arrowhead). (C) SEM image of oviprovector scales membrane (indicated by an arrowhead). (C’) Enlarged image of the rectangular area in (C). Scale bar indicates 100 μm in (C) and 10 μm in (C’).

To elucidate the changes in the oviprovector scale arrangement, the ovipositors of the two species and their closely related species, *D. biarmipes*, and an outgroup, *D. melanogaster*, were dissected and mounted to visualize the scales (Fig. 2A–E, Fig. S1). The degree of sclerotization and size of the scales were variable among these species. The oviprovector scales in *D. suzukii* and *D. subpulchrella* were more sclerotized than those in *D. melanogaster* and *D. biarmipes* (Fig. 2B–E). The size of the scales measured by lateral area of the protrusion was ~1.6–2.1 times larger in *D. suzukii* compared to *D. melanogaster* (Fig. 2F). The size was ~0.47 times smaller in *D. biarmipes* than in *D. melanogaster*. Between *D. suzukii* and *D. subpulchrella*, the scale size was ~1.8–2.3 times larger in *D. subpulchrella* than in *D. suzukii*. These measurements indicate that the sclerotization and the drastic enlargement of the scale are likely to have taken place in the common ancestor of *D. suzukii* and *D. subpulchrella* after the split from *D. biarmipes* along with the elongation of the ovipositor plates. Further divergence in size occurred between the lineages leading to *D. suzukii* and *D. subpulchrella*.

**Fig. 2.**
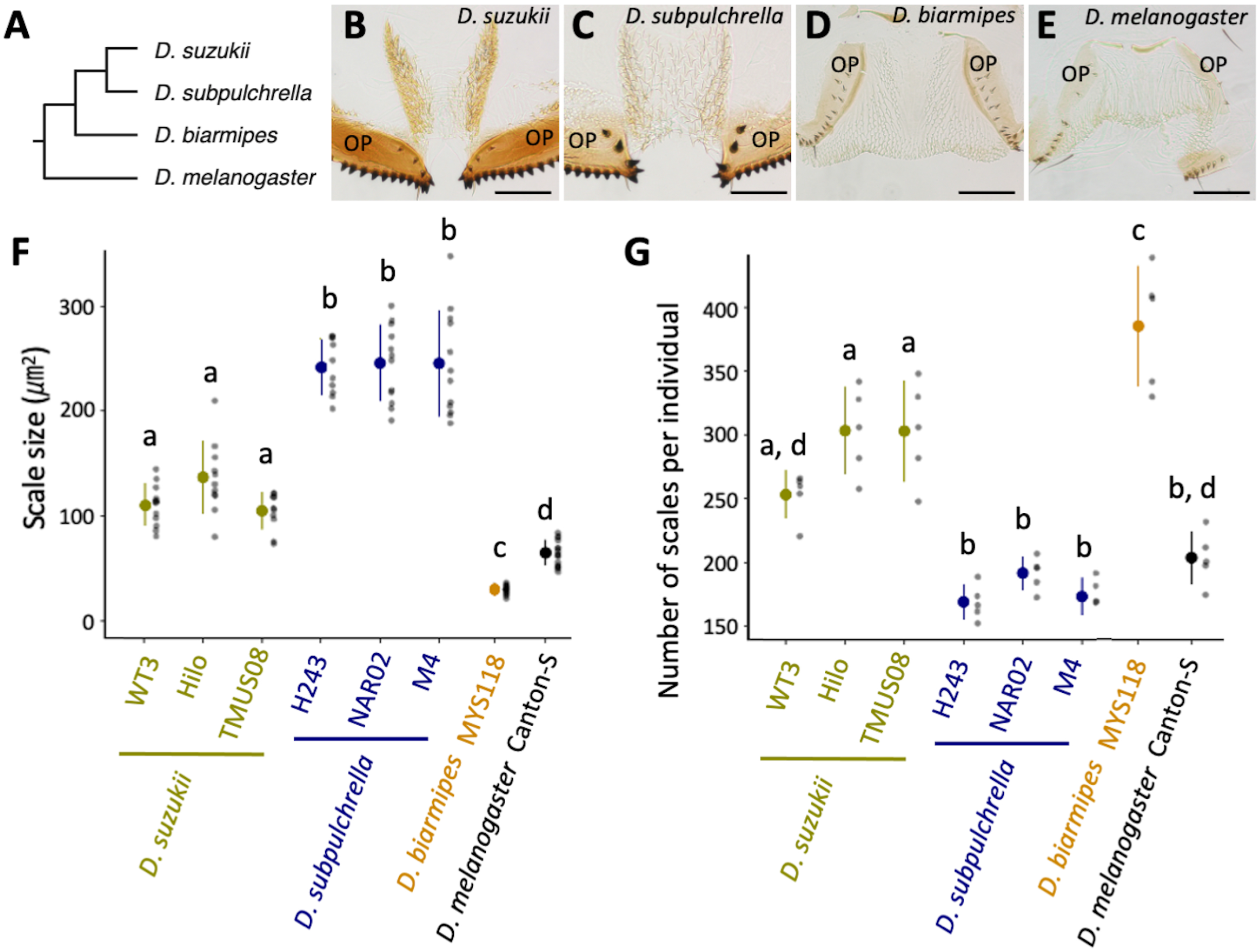
Diversification of oviprovector scales in *Drosophila*. (A) Phylogenetic relationship of species used in this study. Tree topology is based on Suvorov et al. [57]. (B–E) Images of oviprovector scales from dissected ovipositors in different species. Ovipositor plates are indicated as OP. Sale bar indicates 100 μm. (F) Scale size measured from ten individuals from each strain. (G) Number of scales per individual counted in five individuals from each strain. Different letters indicate significant differences between strains after Tukey’s multiple comparisons (P < 0.05). Mean and standard deviation are shown as dot and error bar, respectively.

The distribution of the scales on the oviprovector inner surface was also variable. The locations of the scales were confined to the narrow area at the distal opening of the oviprovector in *D. melanogaster* (Fig. 2E). In contrast, the scales were distributed in a broader area toward the proximal end in the other three species (Fig. 2B–D). Comparing *D. suzukii* and *D. subpulchrella*, the former showed a polarized distribution of the scales on the lateral surface of the duct (Fig. 2B), which contrasts with the uniform distribution in the latter species (Fig. 2C). The observation of the oviprovector scales in three additional strains from *D. suzukii* and *D. subpulchrella* indicated that these morphological characteristics are fixed within these species (Additional file 1: Fig. S1). The number of scales was also different among the species, largely reflecting their distribution on the oviprovector membrane (Fig. 2G). These observations suggest that the oviprovector scale arrangement is a rapidly diversifying trait of the female genitalia in *Drosophila*.

### Oviprovector scales are trichomes protruded from single cells expressing *svb*

The shape of the oviprovector scales resembles that of the trichomes on the larval cuticle, which are actin-rich apical extensions from the epidermal cells. To confirm that the oviprovector scales are single-cell projections like other trichomes, the immunostaining of E-cadherin combined with the rhodamine phalloidin (actin) staining was conducted on the developing ovipositor of *D. melanogaster* at 44 and 48 h APF. The confocal images depicted that actin projections are indeed derived from single cells (Fig. 3A, B, B’). The staining using the ovipositors of *D. suzukii* and *D. subpulchrella* at 45 h APF also showed the same structure (Fig. 3E, F). Therefore, the numbers and sizes of the oviprovector scales can be used to infer those features of trichome-forming epidermal cells of the oviprovector membrane. Furthermore, the intensity of phalloidin in the ovipositor at 44 h APF was lower than that at 48 h APF in *D. melanogaster* (Fig. 3A, B). This result indicated that the actin accumulates between these time points and forms scales on the oviprovector membrane.

**Fig. 3.**
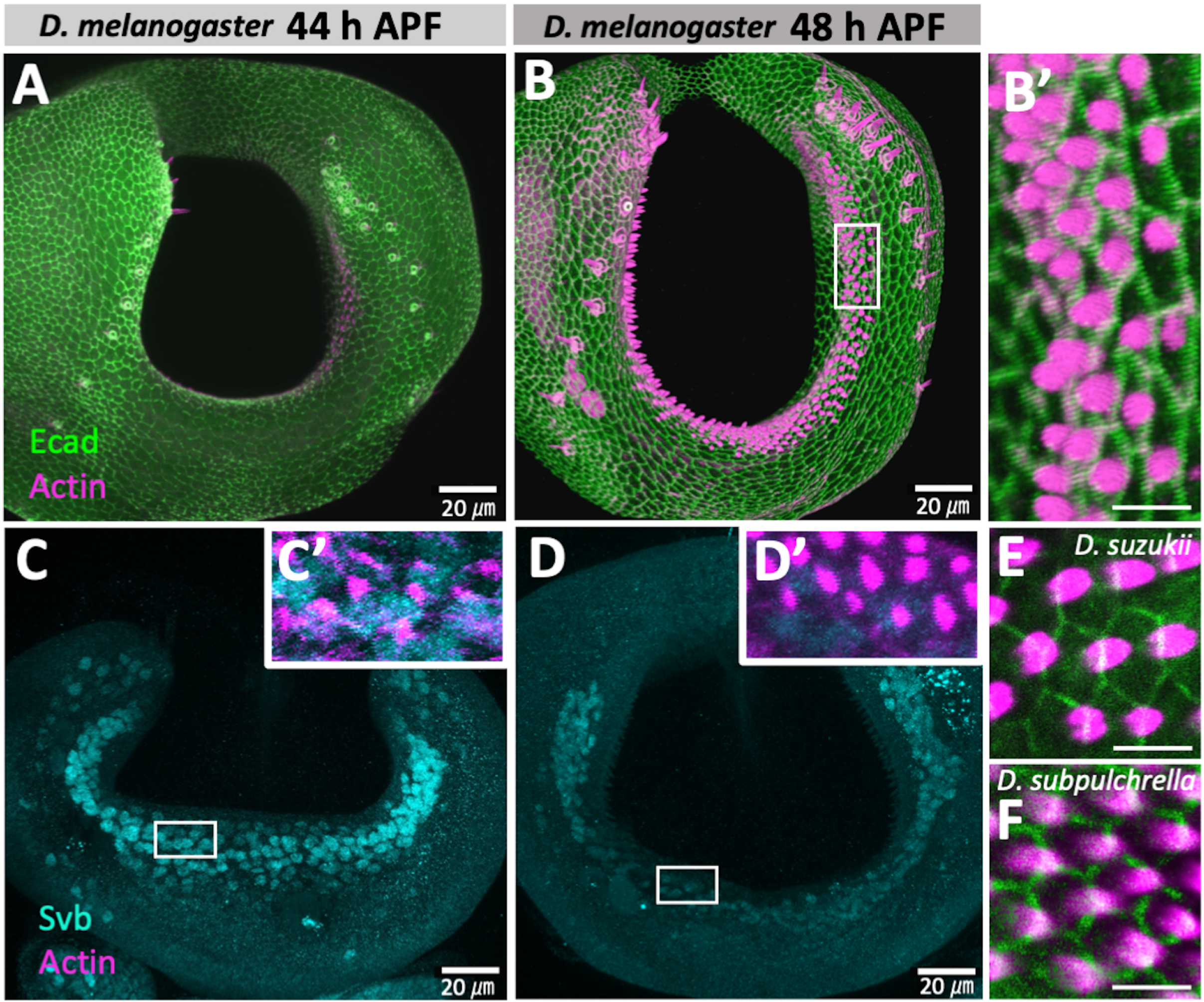
Confocal images of developing ovipositors in *D. melanogaster* (*y, w*), *D. suzukii* (TMUS08), and *D. subpulchrella* (H243). (A) Configuration of E-cadherin (green) and F-actin/phalloidin (magenta) at 44 h APF in *D. melanogaster*. (B) Configuration of E-cadherin (green) and F-actin/phalloidin (magenta) at 48 h APF in *D. melanogaster*. (B’) Enlarged image of the rectangular area in (B). (C) Immunostaining of Svb (light blue) at 44 h APF in *D. melanogaster*. (C’) Enlarged image of the rectangular area in (C) with F-actin/phalloidin (magenta). (D) Immunostaining of Svb (light blue) at 48 h APF in *D. melanogaster*. (D’) Enlarged image of the rectangular area in (D) with F-actin/phalloidin (magenta). (E) Configuration of E-cadherin (green) and F-actin/phalloidin (magenta) in *D. suzukii* at 45 h APF. (F) Configuration of E-cadherin (green) and F-actin/phalloidin (magenta) in *D. subpulchrella* at 45 h APF. Scale bars indicate 20 μm in (A–D) and 5 μm in (B’), (E), and (F).

The gene regulatory network that controls trichome formation is well-characterized in *D. melanogaster*, and *svb* is known to be the master control gene regulating the network [17, 18, 23]. The antibody staining of Svb at 44 and 48 h APF in *D. melanogaster* revealed that this gene product is present in the cells on the inner surface of the developing ovipositor where actin bundles are forming (Fig. 3C, C’, D, D’). This result implied that the trichome gene network is activated in those scale-producing cells.

### Transcriptomic analysis of 48 h APF female genitalia from *D. suzukii* and *D. subpulchrella*

Motivated by the presence of Svb in the cells on the inner surface of the developing ovipositor in *D. melanogaster*, we next asked 1) if the known genes in the network are expressed in the developing female genitalia of *D. suzukii* and *D. subpulchrella*, and 2) if any of the trichome-related genes contribute to the morphological differences of the oviprovector scales between these species. The ovipositor morphogenesis takes place at early pupal stages in *Drosophila* at around 30–54 h APF [21]. A time point was taken at 48 h APF when the projection of the ovipositor plates has proceeded to the mid-step. The extracted RNA samples from the frozen genital tissues of pupae at 48 h APF were subjected to RNA-seq analysis.

The transcripts of 8,666 genes with more than 1 transcript per million (TPM) were detected by mapping the reads to the reference coding sequences (CDS) generated by reciprocal re-annotation (Additional file 1: Table S1, Table S2). Among them, 1,573 genes were differentially expressed between *D. suzukii* and *D. subpulchrella* (false discovery rate (FDR) < 0.01) (Additional file 1: Table S2). Among the homologs of 50 genes that are known to be involved in trichome pattern formation in *D. melanogaster* [18], 46 genes were included in the reference CDS, and 43 genes showed detectable gene expression levels (> 1 TPM) in either of the two species, suggesting the involvement of the genes in the trichome regulatory network. Eleven genes showed differential expression at FDR < 0.01 (Table 1). Five genes, *Cuticular protein 11A* (*Cpr11A*), *krotzkopf verkehrt* (*kkv*), *CG10175*, *CG15005* and *Osiris 24* (*Osi24*), showed higher expression in *D. suzukii*, and six genes, *shavenoid* (*sha*), *morpheyus* (*mey*), *nyobe* (*nyo*), *neyo* (*neo*), *CG31559* and *Reduction in Cnn dots 6* (*Rcd6*), showed higher expression in *D. subpulchrella* (at FDR < 0.01). The master control gene of the trichome regulatory network, *shavenbaby* (*svb*), did not show differential expression between these species at this stage.

**Table 1.** Expression differences of trichome related genes between *D. suzukii* and *D. subpulchrella*.

| Gene name | log<sub>2</sub>CPM | Log<sub>2</sub>Fold-change<br>(*D. suzukii*/*D. subpulchrella*) | FDR | Function in trichome<br>formation |
| --- | --- | --- | --- | --- |
| *wg* | 5.8 | -0.51 | 5.2×10<sup>-2</sup> | Upstream genes of *svb* |
| *abd-A* | 7.2 | -0.20 | 3.2×10<sup>-1</sup> | Upstream genes of *svb* |
| *Ubx* | 2.3 | -0.61 | 3.8×10<sup>-1</sup> | Upstream genes of *svb* |
| *hh* | 4.9 | -0.28 | 3.6×10<sup>-1</sup> | Upstream genes of *svb* |
| *lin* | 5.1 | -0.14 | 6.2×10<sup>-1</sup> | Upstream genes of *svb* |
| *Ser* | 7.8 | -0.43 | 2.0×10<sup>-2</sup> | Upstream genes of *svb* |
| *rho* | 4.0 | 0.24 | 5.8×10<sup>-1</sup> | Upstream genes of *svb* |
| *aos* | 4.0 | -0.49 | 1.2×10<sup>-1</sup> | Upstream genes of *svb* |
| *SoxN* | low expression<sup>a</sup> | - | - | Upstream genes of *svb* |
| *D* | 3.2 | -0.37 | 4.3×10<sup>-1</sup> | Upstream genes of *svb* |
| *tal* | n.a.<sup>b</sup> | - | - | Upstream genes of *svb* |
| *ovo/svb* | 5.9 | -0.43 | 3.0×10<sup>-1</sup> | Master control gene |
| <u>*sha*</u> | <u>8.5</u> | <u>-0.78</u> | <u>7.6×10<sup>-5</sup></u> | Actin bundling |
| *sn* | 8.8 | -0.02 | 9.2×10<sup>-1</sup> | Actin bundling |
| *f* | 9.6 | 0.02 | 9.4×10<sup>-1</sup> | Actin bundling |
| *wasp* | 5.8 | 0.23 | 3.2×10<sup>-1</sup> | Actin bundling |
| *mwh* | 8.8 | -0.51 | 6.7×10<sup>-2</sup> | Actin bundling |
| *Actn* | 10.0 | 0.17 | $4.0 \times 10^{-1}$ | Actin bundling |
| *y* | 4.4 | -0.76 | $2.9 \times 10^{-2}$ | Cuticle |
| *ChLD3* | 5.5 | 0.43 | $2.0 \times 10^{-1}$ | Cuticle |
| *Cda5* | 6.0 | 0.40 | $2.1 \times 10^{-1}$ | Cuticle |
| *Cpr11A* | <u>3.0</u> | <u>1.34</u> | <u>$3.1 \times 10^{-4}$</u> | Cuticle |
| *kkv* | <u>9.7</u> | <u>0.92</u> | <u>$2.1 \times 10^{-4}$</u> | Cuticle |
| *Peritrophin-A* | low expression<sup>a</sup> | - | - | Cuticle |
| *m* | 10.6 | -0.14 | $6.5 \times 10^{-1}$ | Membrane matrix |
| *dyl* | 9.8 | 0.74 | $9.9 \times 10^{-2}$ | Membrane matrix |
| *mey* | <u>11.7</u> | <u>-0.54</u> | <u>$6.3 \times 10^{-3}$</u> | Membrane matrix |
| *nyo* | <u>10.3</u> | <u>-0.63</u> | <u>$2.0 \times 10^{-3}$</u> | Membrane matrix |
| *tyn* | 12.7 | -0.13 | $6.4 \times 10^{-1}$ | Membrane matrix |
| *neo* | <u>12.2</u> | <u>-0.53</u> | <u>$7.7 \times 10^{-3}$</u> | Membrane matrix |
| *zye* | 5.4 | 0.20 | $5.9 \times 10^{-1}$ | Membrane matrix |
| *cyr* | 8.6 | -0.07 | $8.7 \times 10^{-1}$ | Membrane matrix |
| *CG16798* | 7.9 | -0.18 | $5.5 \times 10^{-1}$ | Membrane matrix |
| *CG12814* | 7.2 | -0.40 | $5.7 \times 10^{-2}$ | Membrane matrix |
| *CG4702* | 11.0 | 0.14 | $6.6 \times 10^{-1}$ | Unknown |
| *CG8303* | 5.7 | -0.04 | $9.2 \times 10^{-1}$ | Unknown |
| <u>CG10175</u> | <u>3.2</u> | <u>1.95</u> | <u>$6.2 \times 10^{-6}$</u> | Unknown |
| *CG14395* | 9.2 | -0.65 | $4.8 \times 10^{-2}$ | Unknown |
| <u>CG15005</u> | <u>4.9</u> | <u>1.61</u> | <u>$4.7 \times 10^{-6}$</u> | Unknown |
| *CG15022* | low expression<sup>a</sup> | - | - | Unknown |
| <u>CG31559</u> | <u>7.2</u> | <u>-0.99</u> | <u>$8.0 \times 10^{-10}$</u> | Unknown |
| *CG32694* | 3.6 | -0.10 | $8.8 \times 10^{-1}$ | Unknown |
| *CG45064* | n.a.<sup>b</sup> | - | - | Unknown |
| *Sox21b* | n.a.<sup>b</sup> | - | - | Unknown |
| <u>Osi24</u> | <u>6.5</u> | <u>0.98</u> | <u>$3.7 \times 10^{-3}$</u> | Unknown |
| *Spn88Ea* | 7.4 | -0.14 | $5.4 \times 10^{-1}$ | Unknown |
| *Tg* | n.a.<sup>b</sup> | - | - | Unknown |
| <u>Rcd6</u> | <u>4.0</u> | <u>-0.78</u> | <u>$3.3 \times 10^{-3}$</u> | Unknown |
| *PH4aEFB* | 6.0 | -0.01 | $9.7 \times 10^{-1}$ | Unknown |
| *ImpE1* | 12.5 | 0.26 | $2.7 \times 10^{-1}$ | Unknown |

<sup>a</sup> : TPM < 1 in both species
<sup>b</sup> : CDS was not available in our analysis.

### *sha* is expressed in the inner cavity of the developing ovipositor

The trichome network genes were detected by the RNA-seq analysis. To confirm that the network pathway is activated in the developing oviprovector of *D. suzukii* and *D. subpulchrella*, an in situ hybridization of an effector gene, *sha*, was conducted to examine its expression at the inner cells of the developing ovipositors. *sha*, is a direct target of Svb and functions in bundling actin fibers on the apical surface of the cells [16, 24]. The gene also showed a differential expression level between the two species (Table 1).

At 44 and 48 h APF, when the duct inside the ovipositor plates is still not closed at the distal end, a clear signal was detected at the inner cavity of the ovipositors in both *D. suzukii* and *D. subpulchrella* (Fig. 4A–E). While the expression domain was split into two lateral areas on the inner surface of the cavity in *D. suzukii* (Fig. 4B, D), only one domain connected at the ventral side of the cavity was observed in *D. subpulchrella* (Fig. 4C, E). The pattern resembled that of the adult oviprovector scale distributions of the two species (Fig. 2B, C), indicating that this gene in the trichome regulatory network, is expressed in the cells where trichomes develop at a later stage in these species.

**Fig. 4.**
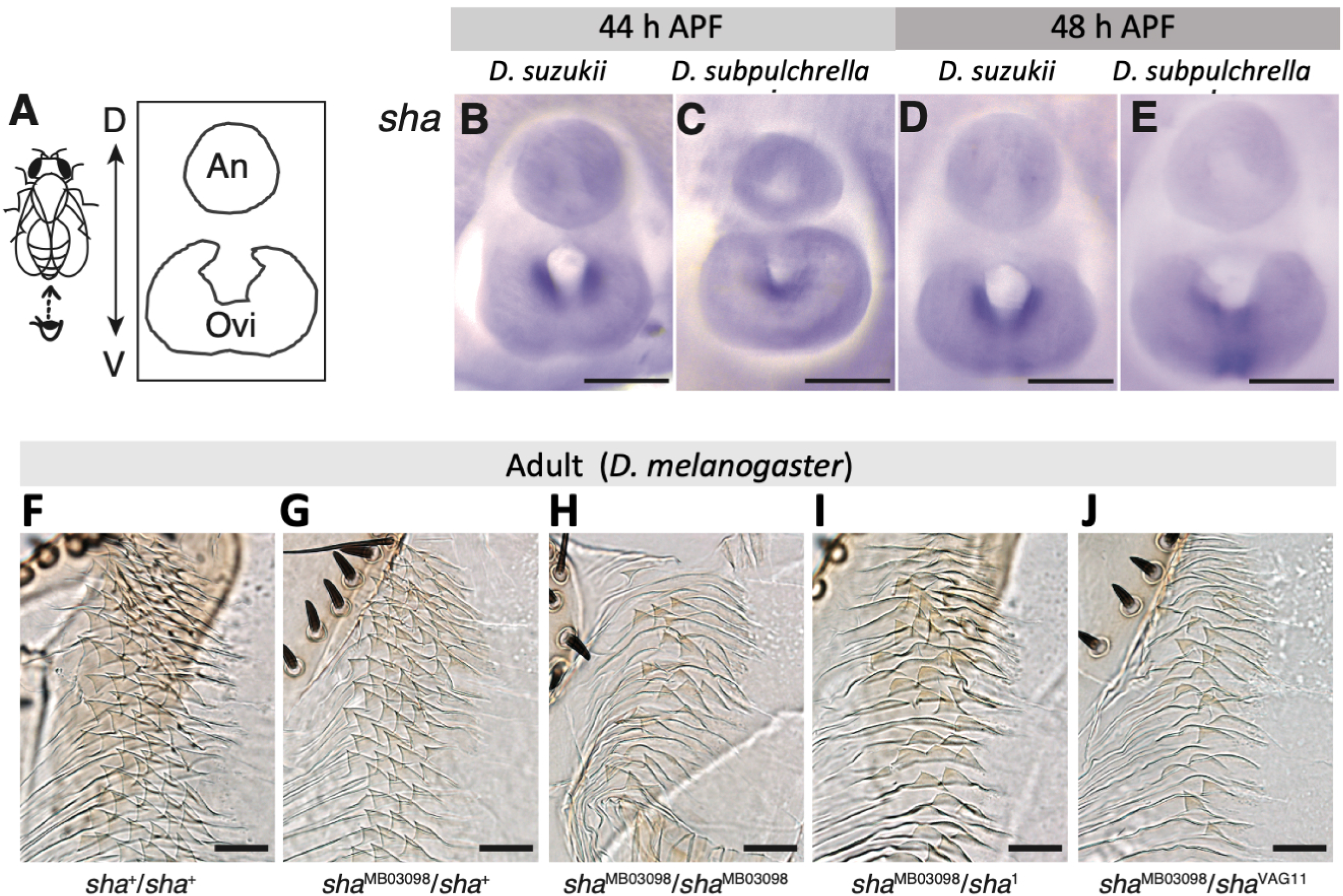
*sha* expression in the developing ovipositor and the oviprovector scale defects in sha mutants. (A) Schematic image of the developing female terminalia. Abbreviations indicate analia (An) and ovipositor (Ovi). (B) in situ hybridization of *sha* at 44 h APF in *D. suzukii*. (C) in situ hybridization of *sha* at 44 h APF in *D. subpulchrella*. (D) in situ hybridization of *sha* at 48 h APF in *D. suzukii*. (E) in situ hybridization of *sha* at 48 h APF in *D. subpulchrella*. (F) Oviprovector scales of Canton-S (*sha*^+^/*sha*^+^). (G) Oviprovector scales of *sha*^MB03098^/*sha*^+^. (H) Oviprovector scales of *sha*^MB03098^/*sha*^MB03098^. (I) Oviprovector scales of *sha*^MB03098^/*sha*^1^. (I) Oviprovector scales of *sha*^MB03098^/*sha*^VAG11^. Scale bars indicate 100 μm in (B–E) and 20 μm in (F–J).

### Oviprovector scales of *D. melanogaster sha* mutants

To further investigate whether *sha* is involved in the development of trichomes on the oviprovector, mutant phenotypes were observed in *D. melanogaster*. In the wild type *D. melanogaster*, the oviprovector scales (trichomes) are triangular with hook-like tips and highly arrayed configuration (Fig. 2E, 4F, Additional file 1: Fig. S2A). The scales of a homozygous *sha* mutant, *sha*^MB03098^, showed a malformation lacking most of the spikes (Fig. 4H, Additional file 1: Fig. S2B). Similar defects were observed in two other *sha* mutant alleles that were tested in heterozygotes, *sha*^1^/*sha*^MB03098^ and *sha*^VAG11^/*sha*^MB03098^ (Fig. 4I, J). Mutants of the genes involved in the apical extracellular matrix formation, *trinity* (*tyn*) and *miniature* (*m*) [17, 25], showed a similarly arrayed but slightly smaller trichomes compared to the wild type (Additional file 1: Fig. S2C, D). These results corroborate that the gene regulatory network underlying the development of larval denticles is likely to be adopted to the developmental system of oviprovector scales.

## Discussion

The oviprovector scales in *D. suzukii* and *D. subpulchrella* were distinct in size and distribution from those of *D. biarmipes* and *D. melanogaster* (Fig. 2) as anticipated from their uniquely enlarged ovipositor plates [8, 20, 21]. *D. suzukii* and *D. subpulchrella* have shifted their oviposition sites from rotting to ripening fruits, and the shift has promoted the evolution in various traits including the ovipositor morphology as well as the mechanosensory and chemosensory perception [8, 20, 21, 26–30]. The ovipositor is an essential device for the flies to deposit eggs to the appropriate substrate; therefore, the changes in ovipositor shape should accompany an adjustment of oviprovector morphology to support egg-laying effectively. The exact function has not been explored in detail. However, the arrayed trichomes tilting towards the distal direction might effectively reduce the friction between the surfaces of the egg and the duct when the egg is moving toward the distal opening. Alternatively, they might be functioning as small hooks to grab and push the egg toward the vulva opening by reversing the membrane [14, 22]. Since similar oviprovector scales are present in other insect species (e.g., [14, 31–33]), they must share a common functional role in oviposition.

The ovipositor plates of *D. suzukii* are less curved and become narrower at the distal tip than those of *D. subpulchrella* [8, 20]. The change in shape could have affected the mechanics of egg deposition and provoked a change in the distribution of scales on the oviprovector. We speculate that applying forces from the lateral direction by a pair of plates to push the eggs along the longitudinal direction of the ovipositor required the alignment of scales on the lateral surface of the oviprovector in *D. suzukii*. Whereas due to the curved ovipositor plates of *D. subpulchrella*, the egg might be slightly pushed towards the ventral side of the ovipositor. Thus, the scales on the ventral surface of the oviprovector might effectively reduce the surface friction or grab the egg by the spiny tips. These speculations require rigorous validation by a detailed biophysical analysis. Nevertheless, such fine-tuning of the egg transporting mechanics upon the changes in external morphology of the ovipositors might have enforced frequent rearrangements of the oviprovector scales representing a rapidly diversifying morphological trait of the female reproductive tract.

The gene regulatory network underlying trichome development is shared in many body regions, such as in the denticle belts of the larval cuticle and on the surface of the wings and adult cuticle (reviewed in [18, 34]). We have shown by the immunostaining of Svb and the RNA-seq analysis using the developing female genitalia that the *svb*-regulated pathway is likely to be involved in the formation of the trichomes on the oviprovector membrane. The expression patterns of *sha*, a direct target of Svb, at 44 and 48 h APF in *D. suzukii* and *D. subpulchrella* resembled the corresponding patterns of the trichome distribution on the adult oviprovectors of these species (Fig. 4A–E). It is yet to be pursued whether the causal mutations of the interspecific differences in the ovipositor scale characteristics reside in the *cis*-regulatory regions of the effector gene *sha* or those that change the expression patterns of *svb* or other upstream genes.

Some components of the trichome gene regulatory network differ between the larval and leg trichomes [35]. In contrary to various factors regulating *svb* in the embryonic and larval cuticle, the development of trichomes is suppressed by *microRNA-92a* in a cuticular patch, “naked valley”, in the leg (T2 femurs) by regulating the translation of *sha* [36]. In our analysis, it was intriguing to see that *svb* did not show differential expression between the two species despite the significant expression level differences (1.7-fold lower in *D. suzukii*) of *sha* in the developing female genitalia (Table 1). Thus, a possibility that *sha* is directly regulated by microRNAs independently from Svb remains to be investigated. *mey*, *nyo*, and *neo* also showed significantly lower expression (1.4–1.5-fold lower) in *D. suzukii* than in *D. subpulchrella* (Table 1). These genes are suggested to act together to form apical extracellular matrix during the development of the larval denticles [25], therefore, they may be differentially regulated simultaneously in the developing oviprovector scales between the two species. Nevertheless, it should be noted that per nucleus transcript abundance cannot be accurately compared in our data because the number of trichome-producing cells in the oviprovector are different between the two species (Fig. 2G). Further investigation of the detailed expression domains of these genes may reveal some differences in the gene regulatory network components between the oviprovector and other parts of the fly body.

During the ovipositor morphogenesis at the pupal stage, Green et al. [21] have analyzed the size and number of the external cell layer of the ovipositor plate. They concluded that an accelerated cell size expansion instead of an enhanced cell division causes the enlargement of the ovipositors in *D. suzukii*. Since the trichomes on the oviprovector were shown to be single-cell extensions as in the cuticle trichomes (Fig. 3), the differences in the size of the trichomes reflect the differences in the apical cell area. Therefore, the size differences of the trichomes can be utilized to infer cell size differences between species.

The lateral area of the scale in *D. suzukii* was roughly two times larger than that of *D. melanogaster* but about two times smaller than that in *D. subpulchrella*. These size differences imply that the expansion of apical cell areas in *D. suzukii* is not confined to the external cells of the ovipositor plate but also at the inner membrane of the oviprovector, at least in the scale producing cells. Also, the comparisons suggest that the apical cell areas of the scale-producing cells of *D. subpulchrella* expanded more extensively, which might also be the case in the ovipositor plate cells that have not yet been investigated in this species.

It is also interesting to note that the arrayed alignment and the direction of the actin bundles in the developing ovipositors imply planar cell polarity. Mechanisms underlying such polarity are well-studied in other actin-based external bristles and hairs on the adult cuticle and wing (reviewed in [37, 38]) as well as in the multiciliated cells of the mammalian oviduct [39–41]. Further analyses of the changes in cell size and polarity during morphogenesis may uncover the details of the cell dynamics underlying differential degrees of the oviprovector rearrangement upon the ecological niche exploitation in these species.

## Conclusions

The oviprovector scales inside the recently enlarged ovipositors of *D. suzukii* and *D. subpulchrella* were distinct in size and distribution pattern compared to those in a closely related species, *D. biarmipes*, and an outgroup species, *D. melanogaster*. The size of the scales indicated that the apical cell area of the oviprovector has expanded upon the elongation of the ovipositor plates in the two species. Our transcriptome analysis revealed that most of the known genes in the trichome gene regulatory network underlying larval denticle formation are expressed in the developing female genitalia of *D. suzukii* and *D. subpulchrella*. We confirmed the presence of Svb in the inner cavity of the developing ovipositors of *D. melanogaster* and the *sha* expression in the similar regions of *D. suzukii* and *D. subpulchrella*, suggesting that the network is adopted to develop the ovipositor trichomes. Taken together, in parallel to the well-documented cases of rapidly evolving genital structures in *Drosophila* males (e.g., [42–45]), our study opens an opportunity to explore the evolution of a rapidly diversifying morphological trait of the female genitalia. Our study also highlights the gene regulatory network of a functionally distinct and divergent trichome system associated with ecological niche exploitation.

## Methods

### Fly strains

Wild-derived strains of *D. suzukii* (TMUS08, TMUS09, TMUS18, WT3, and Hilo) and *D. subpulchrella* (H243, M4, NAR02, and NAR07) were maintained at 20°C unless stated otherwise. A wild-derived strain of *D. biarmipes* (MYS118) was maintained at 25°C. TMUS08, TMUS09, and TMUS18 were collected in Hachioji, Japan, in 2015. Hilo was collected in Hilo, Island of Hawai’i, USA, in 2017. WT3 strain was obtained from the *Drosophila* Species Stock Center. H243 strain was collected in Hiratsuka, Japan, in 1979. M4 was collected in Matsumoto, Japan, in 1982. NAR02 and NAR07 were collected in Narusawa-mura, Japan, in 2016. MYS118 was collected in Mysore, India, in 1981. *D. melanogaster* Canton-S, *yw* (#1495), and other mutant strains were maintained at 25°C. *sha*^1^ (#107623) and *m*^1^ (#105713) were obtained from KYOTO Stock Center (DGRC). s*ha*^VAG11^ (#32097), *sha*^MB03098^ (#24445), and *tyn*^EP1421^ (#10277) were obtained from Bloomington *Drosophila* Stock Center. All strains were kept under a 12 h light:12 h dark light cycle.

### Observation of ovipositor scale and morphological measurement

Adult female flies (7–14 days after eclosion) were collected and stored in 70% ethanol before dissection. Dissected ovipositors were mounted in 50% Hoyer’s medium and placed at 68°C overnight. Images were captured using an Olympus IX71 inverted microscope equipped with a DP73 camera (Olympus) at ×200 and ×640 magnifications for the entire distribution and single scale measurement, respectively.

Scanning electron micrographs (SEMs) were taken by a JEOL JSM-6510 scanning electron microscope after being coated with gold using a Hitachi E101 ion sputter (Hitachi Ltd., Tokyo, Japan). For this, the oviprovector was everted by gently pushing the abdomen of virgin female samples with fine forceps in absolute ethanol. The ethanol was then substituted with t-butanol, and the specimens were sublimation dried.

The number of ovipositor scales was manually counted under Olympus IX71 inverted microscope. To measure the size, a single scale on the lateral side of the oviprovector membrane was chosen randomly from each dissected image. The size was measured using Fiji version 2.3 [46] and the statistical tests were conducted by using R version 4.1 [47].

### RNA extraction and sequencing

Three independent biological replicates of RNA-seq libraries for TMUS08 and H243 were generated from abdominal tip tissue dissected from 20 female pupae at 48±1 hours after puparium formation (h APF). Female files were collected at the white pupal stage by sorting gonad size and placed in a humid chamber at 25°C. Staged pupae were flash-frozen by placing them on a cooled metal plate with a cake of dry ice. The posterior tip of the abdomen, including the entire developing ovipositor, was cut at the A6–7 tergite by a blade on the plate. Total RNA was extracted using TRIzol Plus RNA Purification Kit (Life Technologies), and the samples were treated by DNaseI (Invitrogen) to avoid DNA contamination. RNA quality was examined using a High Sensitivity RNA ScreenTape for TapeStation (Agilent Technologies).

Sequencing libraries were generated using the KAPA Stranded mRNA-Seq Kit (KAPA Biosystems) and an Adapter Kit (FastGene) using 200 ng of total RNA. Indexed libraries were sent to Macrogen Japan for sequencing HiSeq X Ten (Illumina), producing 150-bp paired-end reads.

### Differential gene expression analysis

Raw fastq files were adaptor trimmed and quality controlled by fastp ver. 0.20.0 [48] with the following criterion: minimum length 50 bp, Q-score per base ≥ 20, the average Q-score ≥ 30 and maximum number of N bases per read = 1. A reciprocal re-annotation pipeline [49] was applied for generating a total of 13,220 reference CDS from *D. suzukii* WT3 [50] and *D. subpulchrella* RU33 [29] CDS (Additional file3). *D. melanogaster* ortholog gene was assigned from reciprocal best hit to *D. melanogaster* gene products (Flybase: dmel-all-translation-r6.43). After mapping by bowtie2 ver. 2.4.4 [51] the number of reads were counted by samtools ver. 1.10 [52]. Differential gene expression analysis was conducted by edgeR [53]. Differentially expressed genes (DEGs) were detected at the significance level of FDR < 0.01. Genes with TPM < 1 in all three biological replicates in either species were filtered out from the analysis.

### Immunohistochemistry and in situ hybridization

Antibody staining was conducted as in a previous study with minor modification [54]. In brief, newly emerged female pupae were incubated at 25°C and fixed at 45 h APF for *D. suzukii* and *D. subpulchrella* and at 44 and 48 h APF for *D. melanogaster*. Rat anti-E-cadherin 1:200 (DCAD2, DSHB) and donkey anti-rat Alexa 488 1:200 (Thermo Fisher Scientific) antibodies were used. Rhodamine phalloidin 1:200 (Thermo Fisher Scientific) staining was performed with a secondary antibody. The antibody for Shavenbaby (Svb) was a gift from Julia Zeitlinger and was used at 1:10. Images of the developing ovipositors of *D. suzukii* and *D. subpulchrella* were captured by using a C2+ confocal microscope (Nikon) at ×200 magnification. Those of *D. melanogaster* were captured at ×63 on a Leica TCS SP8 confocal microscope. Images were processed by using Fiji version 2.3 [46], MorphoGraphX [55] and Photoshop version 23.1 (Adobe).

For in situ hybridization, newly emerged female pupae were incubated at 25°C for 44 and 48 hours. Staged pupae were collected, fixed, and then proceeded to in situ hybridization as in the previous study with minor modification [56]. A 974-bp s*ha* probe were generated using the following primer set with the addition of T7 linker sequence (GATCACTAATACGACTCACTATAGGG); CAAAGTTTGGCGACCAAGGG (*sha*-fwd), and TCCATCTCCTCCTCGAGTGG (*sha*-rev). Images were captured using an Olympus SZX16 light microscope with a DP73 camera (Olympus) at ×16 magnification.

## Abbreviations

h APF: hour after puparium formation
SEM: Scanning electron micrographs
CDS: coding sequence
DEG: differentially expressed gene
FDR: false discovery rate
TPM: Transcripts per million

## Supplementary Information

**Additional file 1: Figure S1** Oviprovector scales from additional strains. **Figure S2** Oviprovector scales of mutants of trichome-related genes in *D. melanogaster*.

**Additional file 2: Table S1** Mapping results of RNA-seq using *D. suzukii* and *D. subpulchrella* female terminalia at 48 h AFP. **Table S2** Differentially expressed genes between *D. suzukii* and *D. subpulchrella* female terminalia at 48 h AFP.

**Additional file 3:** Reciprocally re-annotated CDS. The .zip folder containing reciprocally reannotated reference CDS used for mapping *D. suzukii* and *D. subpulchrella* RNA-seq reads.

## Acknowledgements

We are grateful to Julia Zeitlinger for providing the Svb antibody. We also thank Koichiro Tamura, KYOTO Stock Center (DGRC), and Bloomington *Drosophila* Stock Center for fly stocks and Sawako Shimizu for technical support.

## Authors’ contributions

KMT, KT, and AT conceived and designed the study. GR and MR analyzed *D. melanogaster* genitalia. YK obtained SEM images. KMT and KT performed RNA-seq. KMT, KT, and AT wrote the manuscript. All authors have read and approved the manuscript. KMT and KT contributed equally to this work.

## Availability of data and materials

Raw fastq files are deposited at DDBJ under the accession number DRA009998 and DRA009999 for *D. suzukii* TMUS08 and *D. subpulchrella* H243, respectively. Fly stocks can be provided upon request. No custom code was used, and all analyses are described in the main text. The authors can provide additional information upon request.

## Declaration

### Ethics approval and consent to participate

No ethics approval or consent to participate were needed for the species used in this study.

### Consent for publication

Not applicable.

