## Supplementary material for "Trichomes on female reproductive tract: rapid diversification and underlying gene regulatory network in *Drosophila suzukii* and its related species": Figure S1, Figure S2

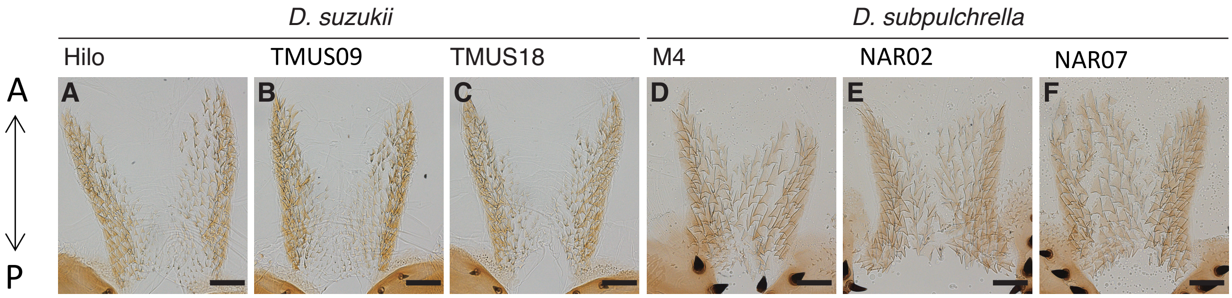


**Fig. S1 Oviprovector scales from additional strains.** (A–C) Oviprovector scales from dissected ovipositors in three additional strains, Hilo, TMUS09, and TMUS18, of *D. suzukii*. (D–F) Oviprovector scales from dissected ovipositors in three additional strains, M4, NAR02, and NAR07 of *D. subpulchrella*. Scale bar indicates 50 μm.

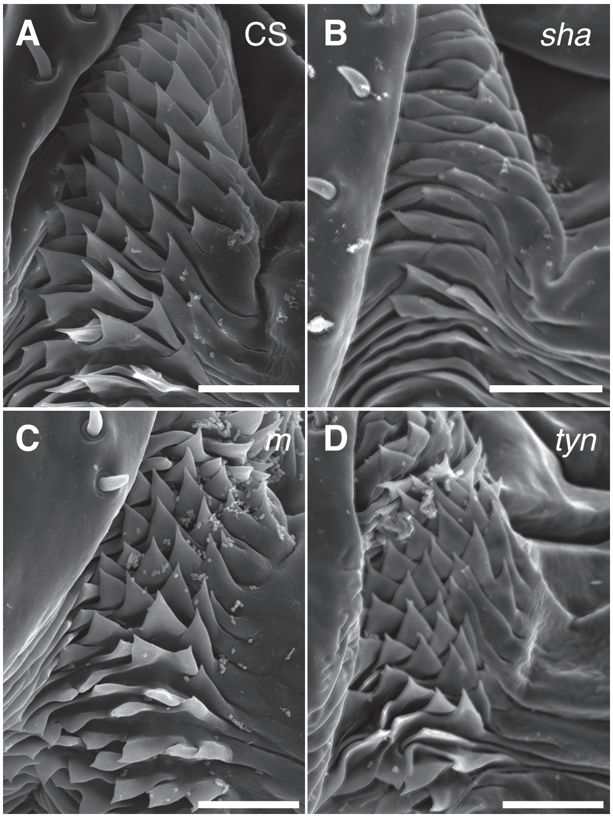


**Fig. S2 Oviprovector scales of mutants of trichome-related genes in *D. melanogaster*.** (A) Wildtype Canton-S. (B–C) Mutant phenotype of trichome patterning genes, *sha* (B), *m* (C) and *tyn* (D). Scale bar indicates 20 μm.
